# Root grooves on two adjacent anterior teeth of *Australopithecus africanus*

**DOI:** 10.1101/222059

**Authors:** Ian Towle, Joel D. Irish, Marina Elliott, Isabelle De Groote

## Abstract

Tooth root grooves and other ante-mortem dental tissue loss not associated with caries found on or near the cementoenamel junction (CEJ) are commonly termed non-carious cervical lesions. Three main processes are implicated in forming these lesions: abrasion, dental erosion, and abfraction. As yet, these lesions have not been described in non-*Homo* hominins. In this study South African fossil hominin collections were examined for evidence of any type of non-carious cervical lesion. Only one individual shows ante-mortem root grooves consistent with non-carious cervical lesions. Two teeth, a mandibular right permanent lateral incisor (STW 270) and canine (STW 213), belonging to the same *Australopithecus africanus* individual, show clear ante-mortem grooves on the labial root surface. These lesions start below the CEJ, extend over a third of the way toward the apex, and taper to a point towards the lingual side. Microscopic examination revealed no clear directional striations. The shape of these grooves is extremely similar to clinical examples of dental erosion, with the lack of striations supporting this diagnosis. These are the oldest hominin examples of non-carious cervical lesions and first described in a genus other than *Homo*; further, the lesions suggest that this individual regularly consumed or processed acidic food items.

## 1. Introduction

Non-carious cervical lesions (NCCLs), resulting from the loss of dental tissue at or near the cementoenamel junction (CEJ), are not uncommon in the dentitions of recent human populations (Levitch et al., 1994; Wood et al., 2008; Grippo et al., 2012). They may result from several processes, the most common of which are abrasion, dental erosion and, to a lesser extent, abfraction (Litonjua et al., 2003; Michael et al., 2009). As such, when NCCLs are present in archaeological and, in this case, fossil hominin remains they may provide some insight into diet and behaviour.

Abrasion is the most common mechanism of NCCLs in archaeological samples; it results from non-masticatory contact of an object with the teeth, and on occasion may relate to cultural or therapeutic behaviour (e.g., Turner and Cacciatore, 1998; Ungar et al., 2001; Novak, 2015; Estalrrich et al., 2016). Dental corrosion, or erosion as it is more commonly called, has been extensively researched in modern (e.g., Zero, 1996; Aubry et al., 2003; Oginni et al., 2003; Grippo et al., 2004) and archaeological human dentitions (e.g., Robb et al., 1991; Indriati and Buikstra, 2001; Ritter et al., 2009; Watson and Haas, 2017). It occurs through the chemical dissolution of dental tissues by acids of non-bacterial origin and can be caused by a range of factors, most notably low pH diets (Indriati and Buikstra, 2001; Oginni et al., 2003; Ritter et al., 2009; Watson and Haas, 2017). Abfraction is the term used to describe tissue breakdown in the cervical region of teeth as a result of extreme occlusal loading from mastication, bruxism, or malocclusion (Lee and Eakle, 1984; Grippo, 1991; Grippo et al., 2012). This process is subject to much debate in the literature, especially in archaeological specimens (Aubry et al., 2003; Michael et al., 2009; Urzúa et al., 2015).

NCCLs have also been described in fossil hominins, the majority of which are ‘toothpick grooves’ (Ubelaker et al., 1969; Boaz and Howell, 1977; Frayer and Russell, 1987; Brown and Molnar, 1990; Milner and Larsen, 1991; Bermudez de Castro et al., 1997; Turner and Cacciatore, 1998; Ungar et al., 2001; Hlusko, 2003; Bouchneb and Maureille, 2004; Kaidonis et al., 2012; Lozano et al., 2013; Tillier et al., 2013; Ricci et al., 2014; Sun et al., 2014; Frayer et al., 2017). Anterior teeth can be affected, but most grooves are present in the interproximal areas of the premolars and molars (Formicola, 1988; Frayer, 1991; Ungar et al., 2001), with buccolingual micro-striations often evident (Bouchneb and Maureille, 2004; Grine et al., 2000; Hlusko, 2003; Lozano et al., 2013; Estalrrich et al., 2016; Sun et al., 2014). The enamel, dentine, and/or cementum are affected depending on groove depth and location, i.e., above, below or directly on the CEJ.

The NCCLs in the above fossil examples are all attributed to abrasion, and are present only in members of the genus *Homo.* Further, no CEJ grooves like those noted above have, to date, been detailed in South African hominins–*Homo* or otherwise (Wallace, 1974; Ungar et al., 2001). For this report, we re-examined the South African collections for evidence of any NCCL type, in specimens assigned to *Paranthropus robustus, Australopithecus africanus, A. sediba, Homo naledi,* and early *Homo.* Of these, one individual is affected, albeit with an unusual form of NCCLs. The aetiology is discussed below and a differential diagnosis conducted.

## 2. Materials and Methods

The teeth in this study include a mandibular right permanent lateral incisor, STW 270, and a mandibular right permanent canine plus several other teeth collectively assigned to STW 213; all are thought to belong to the same individual (Moggi-Cecchi et al., 2006). These specimens originate from Sterkfontein Member 4 and date to 2.8–2.4 Ma (Pickering et al., 2004). Although many consider material from Member 4 to belong to *A. africanus,* others suggest multiple species may be present (e.g., Calcagno et al., 1999; Lockwood and Tobias, 1999; Wood and Richmond, 2000; Moggi-Cecchi, 2003; Grine et al., 2013; Fornai et al., 2015). That said, the specimens studied here are routinely described as *A. africanus.* The only prior mention of the antemortem grooves described here is from Moggi-Cecchi et al. (2006), where they were used to associate STW 270 with 213. Here, the teeth were first examined macroscopically, followed by microscopic observations using a Leica© DMS300 digital microscope (variable magnification ranging from 15 to 40X). Evidence of postmortem damage, other pathology, wear or developmental defect on the teeth were also recorded and are described.

## 3. Results

STW 270 exhibits wear on the incisal edge with a strip of exposed dentine (wear grade 4; Smith, 1984), suggesting that the tooth was in occlusion for some time. An antemortem, concave depression, i.e., shallow groove is evident on the labial root surface just below the CEJ, which extends more than one third of the way toward the apex (Figure 1A). Closer examination revealed no directional antemortem striations (Figure 1B). The groove spreads around to the interproximal areas of the root, narrowing to a point towards the lingual side (Figure 2). The process that created the groove does not appear to have affected other dental tissues, although slight postmortem damage towards the CEJ on the labial surface may mask some evidence. Comparisons with other *A. africanus* teeth show that this is clearly not a morphological feature.

**Figure 1.**
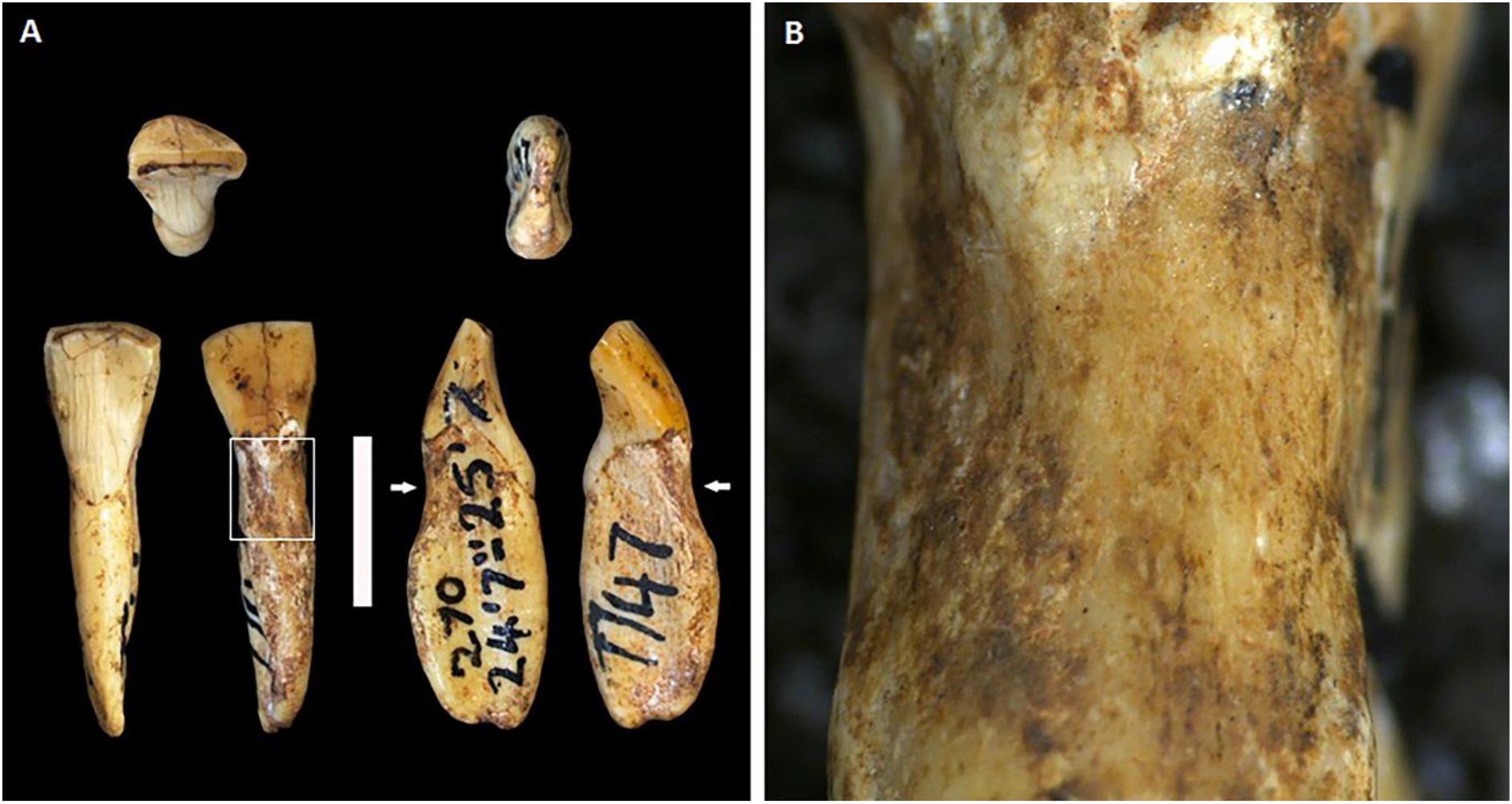
STW 270 (A. *africanus*) right mandibular lateral incisor. A) Bottom row from left to right: lingual, labial, mesial, and distal. White arrows and square highlight the location of the groove. The white bar is 1 cm long. B) Close-up of the groove (white square in A), showing no directional striations and a smooth surface.

**Figure 2.**
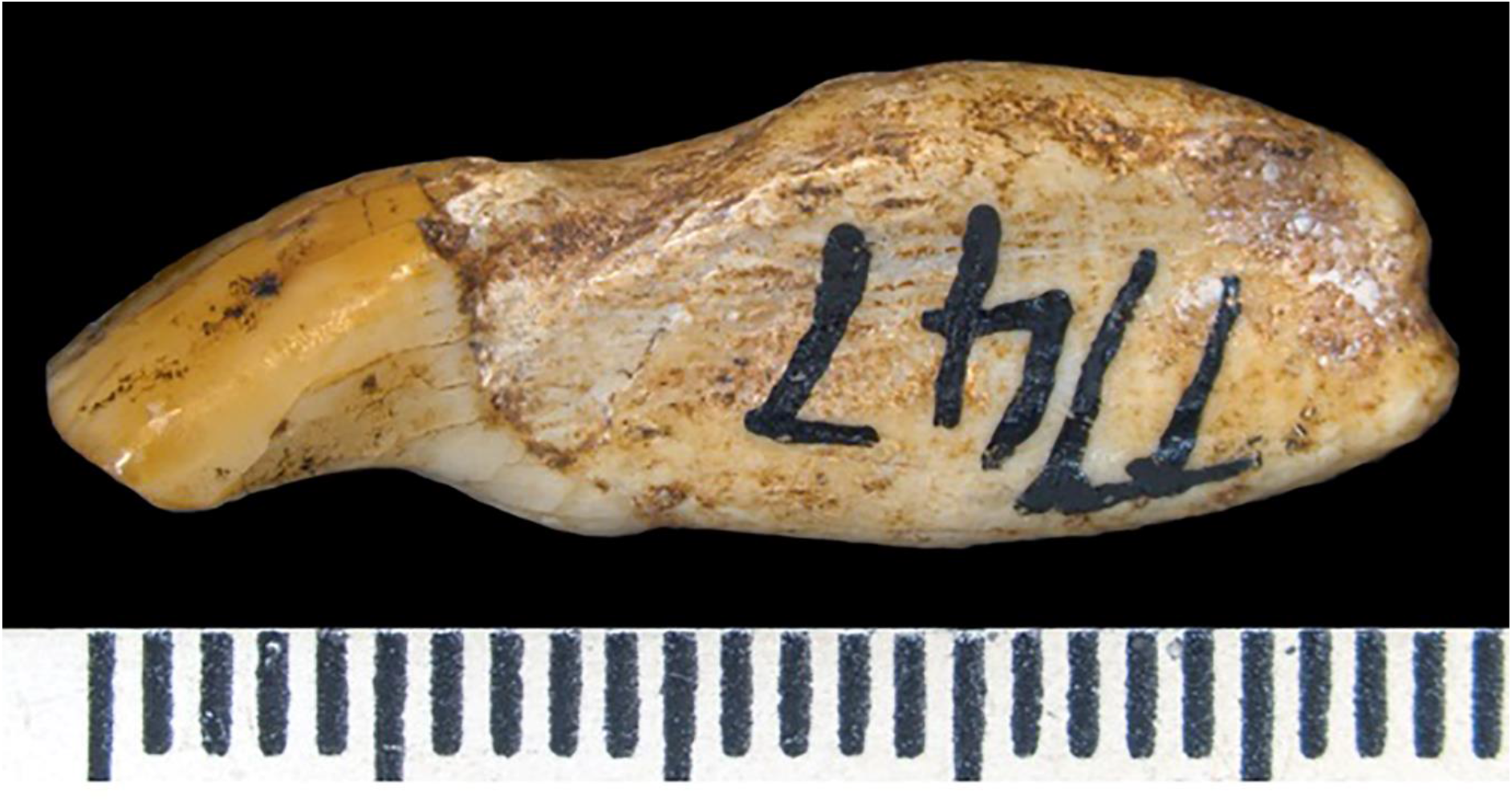
STW 270 (A. *africanus*) right mandibular lateral incisor. Distal side, showing the root groove tapering toward the lingual surface.

A smaller depression/shallow groove is visible in the same position on the root of the adjacent canine, STW 213 (Figure 3). However, this tooth has greater postmortem damage than STW 270 so the full extent of the groove is more difficult to interpret. The shape appears comparable, and grooves in both teeth originate just below the CEJ and extend 5-6 mm down the root. The NCCL on the canine is not as deep as that on the incisor, but given the location and similarity in appearance it is likely that both share a common aetiology.

**Figure 3.**
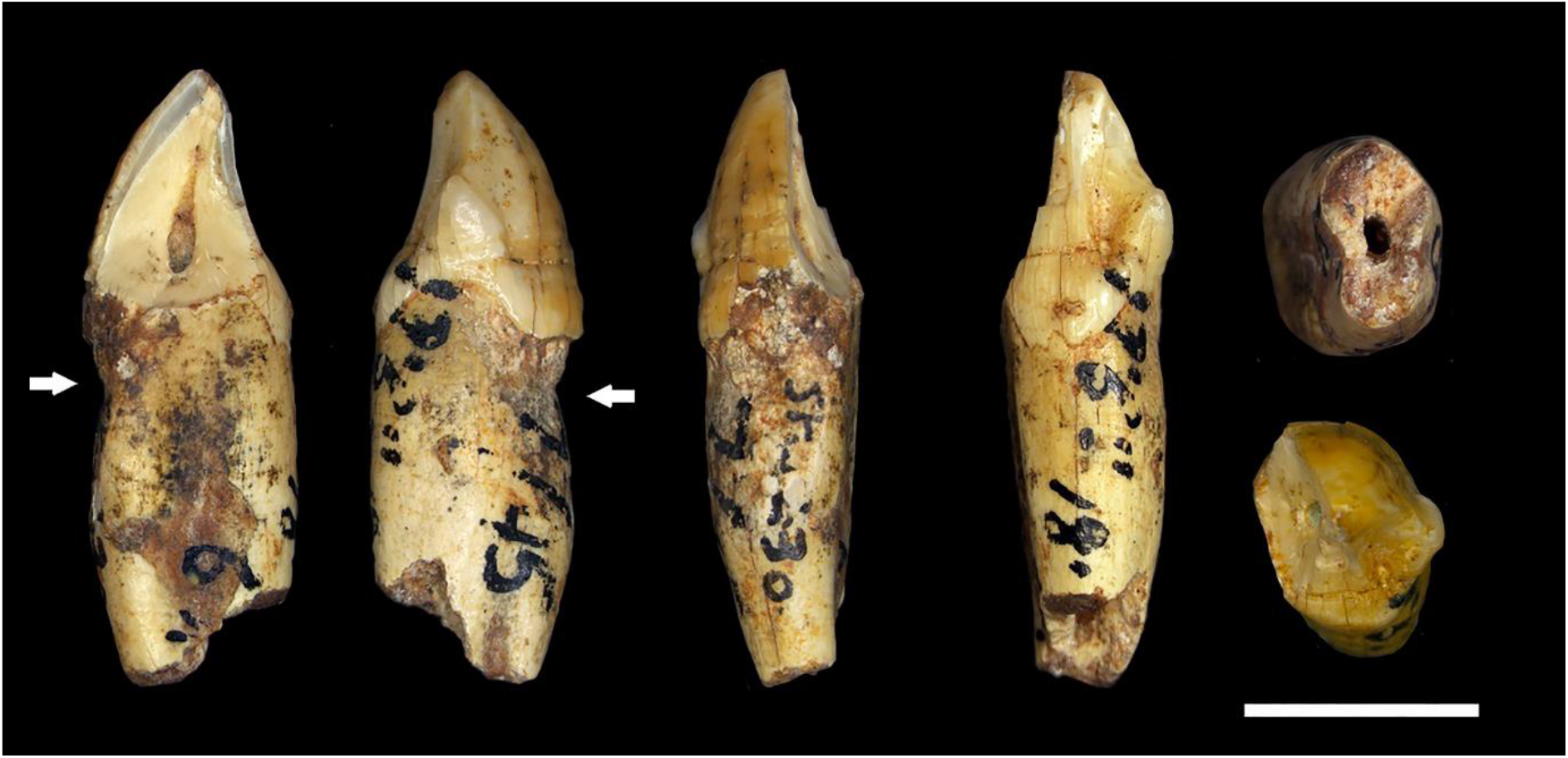
STW 213, mandibular right canine. Left to right: mesial, distal, labial, lingual; upper right: root tip; lower right: crown top.

The other STW 213 teeth do not have root grooves, though only the mandibular first and second molars, premolars and left canine are present. Clear deep furrows are present on the enamel of the right premolars and right canine that are thought to be linear enamel hypoplasia (Guatelli-Steinberg, 2003). The left premolars show no enamel defects. Therefore, the enamel furrows on the right teeth are either not systemic hypoplastic defects or more than one individual is represented in STW 213. There is no evidence of caries or calculus.

## 4. Differential diagnosis

The STW 270 groove appears similar superficially to one described by Novak (2015) in an historic example, i.e., shallow labial groove on a lower canine that extends one third of the way down the root from the CEJ and thought to result from habitual abrasion. As noted, striations are not evident on the *A. africanus* teeth, and the shallow grooves wrap uniformly around both incisor and canine from the labial surface, tapering into the interproximal areas. The latter are also clearly different from typical toothpick grooves associated with specimens of *Homo* (e.g., Frayer and Russell, 1987; Ungar et al., 2001; Lozano et al., 2013; Sun et al., 2014; Tillier et al., 2013; Frayer et al., 2017). Evidence for abfraction-caused NCCLs is limited, but the shallow grooves in STW 270 and 213 do not have the representative wedge shape said to be associated with this process (Michael et al., 2009; Grippo et al., 2012).

Instead, the *A. africanus* grooves best fit an aetiology of erosion, since they are shallow with smooth surfaces (Wood et al., 2008; Levitch et al., 1994). They appear indistinguishable from examples of dental erosion in clinical studies. For example, Bader et al. (1993) highlight a case that also presents a depression/shallow groove on the labial root surface that tapers to a point toward the interproximal areas–along the gingival margins. Additionally, Grippo et al. (2012) describe an upper canine with the erosive lesion extending a significant way down the labial surface of the exposed root. The way in which the shallow groove on the *A. africanus* teeth wrap around the root from the labial surface may suggest they also followed the gingival margins, which is more indicative of erosion. Further, the depressions vary in width but, again, are smooth/uniform as they taper into the interproximal surfaces, which is also suggestive of erosion (Aubry et al., 2003).

If acid was the cause, a gastric origin can be discounted–if not because of the seeming implausibility, then for the reason that lingual tooth surfaces would be affected (Loch et al., 2013). More likely the cause is diet-related. For example, Ritter et al. (2009) report similar NCCLs in archaeological groups that consumed citrus fruits. Another archaeological example described by Watson and Haas (2017; see their Fig. 7) is also akin to that described herein. Shallow labial depressions/grooves are present on some roots of anterior mandibular teeth in a group of prehistoric foragers, though in association with LSAMAT, i.e., lingual surface attrition of the maxillary anterior teeth (Irish and Turner, 1987, 1997). This finding led Watson and Haas (2017) to suggest oral processing of tubers yielded both types of lesions. Tubers are variably acidic, and gingival and alveolar recession may result from stripping and consuming them in a raw state. The present NCCLs could have resulted from such a process, where acidic plant or other food items were ‘processed’ in the mouth, or otherwise just placed in contact with the anterior teeth for long enough periods to erode dentine.

In sum, this is the oldest fossil hominin example of NCCLs, and the first described in a genus other than *Homo.* The shallow root grooves on the STW 270 incisor and 213 canine are identical to examples of dental erosion in the clinical and archaeological literature. Because of their uniqueness, little can be learned about the diet or behaviour of *A. africanus;* however, the grooves do at least attribute some individuality to this particular *A. africanus* individual.

## Acknowledgements

We thank B. Zipfel for access to the collections at the University of the Witwatersrandand and S. Potze from the Ditsong Museum of South Africa for access to their collections. We would also like to thank J. Moggi-Cecchi, D. Guatelli-Steinberg and M. Novak for assistance and advice during this project. This research was supported by a studentship to the first author from Liverpool John Moores University.

